# Identifying genes underlying parallel evolution of stromal resistance to placental and cancer invasion

**DOI:** 10.1101/2025.04.07.647587

**Authors:** Yasir Suhail, Wenqiang Du, Junaid Afzal, Gunter Wagner, Kshitiz

## Abstract

Stromal regulation of cancer dissemination is now well recognized, however causal stromal genes and factors have yet not been identified. Placental invasion into endometrial stroma phenotypically is similar to cancer dissemination. There are also vast differences in the extent of placental invasion across mammals, with human pregnancies at the higher extreme end of invasiveness. We have previously demonstrated that epitheliochorial species, characterized by non-invasive placentation, have acquired stromal resistance to placental invasion, correlating with low cancer malignancy rates. Similarly, decidualization of human endometrial fibroblasts confers resistance to placental invasion. We hypothesized that both trajectories may convergently use similar pathways, using endometrial fibroblasts from 9 species to identify putative ELI-D1 genes underlying stromal vulnerability to invasion. ELI-D1 were negatively enriched in T1-T2 stage transition in many human cancers, typically preceding dissemination, bioinformatically validating their contribution to dissemination-preceding fibroblast activation in cancers. Finally, we have identified candidate transcriptional regulators underlying variation in ELI-D1 genes across eutherians, functionally showing that a key transcriptional regulator of ELI-D1 genes across species, Nr2f6, can regulate stromal resistance to invasion in human fibroblasts. Our comparative approach provides us with a putative, limited, gene-set linked to stromal vulnerability in human cancers, providing therapeutic opportunities to target cancer dissemination.

## Introduction

Placentation substantially varies across mammals, from deeply invasive (hemochorial) to non-invasive (epitheliochorial). This variation is also correlated with differences in rates of cancer malignancy rates in these mammals^1^. We have explained the connection between placental and cancer invasion in an evolutionary framework called **Evolved Levels of Invasibility (ELI)**, showing that endometrial fibroblasts (ESFs) have evolved in epitheliochorial species to resist placental invasion^2^. ELI states that reduced cancer metastasis in epitheliochorial species is a secondary, systematic effect of this phenomenon, likely due to correlated evolution of stroma across other tissues ^3,4^. We have also shown that ELI genes, which are derived from the comparative stromal evolution across epitheliochorial and hemochorial mammalian stroma, are significantly prognostic in predicting survival in human melanoma patients. Identifying the stromal genes that have evolved to confer resistance, or vulnerability to invasion from epithelia, will provide new opportunities to mechanistically understand desmoplastic reaction, as well as develop stroma-targeted therapies to manage cancer metastasis. This objective is however challenging to achieve, because differential gene expression between distant species reveal thousands of genes, most unrelated to any given phenotype. Further, pooled genome-wide gene perturbation libraries are not feasible for an enrichment of a collective phenotype of invasibility.

We therefore identified stromal genes that have changed along with evolution of stromal resistance at the maternal fetal interface with another parallel process at the same site, decidualization of endometrial stroma. Within hemochorial species, ESFs differentiate into a decidual state, which is evolutionarily adapted from the fibroblast activation response to the wounding of endometrial epithelium by implantation, a process similar to the wounding of basal lamina by a growing tumor. Crucially, we have found that decidualization is a differentiation process, which markedly reduces trophoblast invasion by increased matrix production^5^, and actomyosin force generation^6^. Locally, decidualization can therefore be understood as an endometrium specific adaptation to regulate trophoblast invasion.

We therefore identified stromal genes that have changed along with the evolution of epitheliochorial placentation, while belonging to the key decidualization pathways --- based on the assumption that both trajectories may have converged onto similar pathways to achieve stromal resistance. Our approach has identified limited number of high probability genes regulating invasibility, obviating the need for a screen for discovery. This gene-set termed ELI-D1 (ELI and Decidual combined 1) primarily contains negative regulators of cAMP/Protein Kinase A pathway. Interestingly, ELI-D1 was negatively enriched in many TCGA (The Cancer Genome Atlas) cancer genes associated with T1 to T2 stage transition which precedes stromal trespass. Finally, we identified the transcription factor binding sites that had changed in the promoter regions of ELI-D1 genes, correlating their copy numbers with the downstream gene expression. These transcription factors are likely to have contributed to the pattern expression of ELI-D1 genes across mammals. Finally, using a nanopatterned assay to quantitatively measure stromal regulation of invasion, we confirm the most likely transcription regulator of a plurality of ELI-D1 genes, Nr2f6, to be causal in conferring increased vulnerability to invasion in human ESFs.

## Results

### Identifying stromal gene candidates explaining vulnerability to placental invasion in humans

Previously, using RNAseq data from ESFs isolated from 9 eutherian mammals, we identified ELI genes whose expression correlate with placental invasion (**Fig 1A**). We used Accelerated Nanopatterned Stromal Invasion Assay (ANSIA), which we had previously developed to measure stromal resistance (or vulnerability) to invasion^2^ to invasion of HTR8 (human extravillous trophoblasts derived), and F3 (bovine trophoblast derived) into monolayers of human and bovine ESFs (**Fig. 1B-C**). bESFs completely resisted invasion of F3, while hESFs were vulnerable to HTR8 invasion (**Fig. 1C-D**). We similarly confirmed that decidualization increases hESF resistance to HTR8 spheroid invasion on anisotropic monolayers of hESF, undifferentiated or decidualized (dESFs) prior to experimentation (**Fig. 1F-G**). Both the evolution of epitheliochorial placentation, as well as decidualization were associated with large, and significant increase in ESF resistance to corresponding trophoblast invasion.

**Figure 1.**
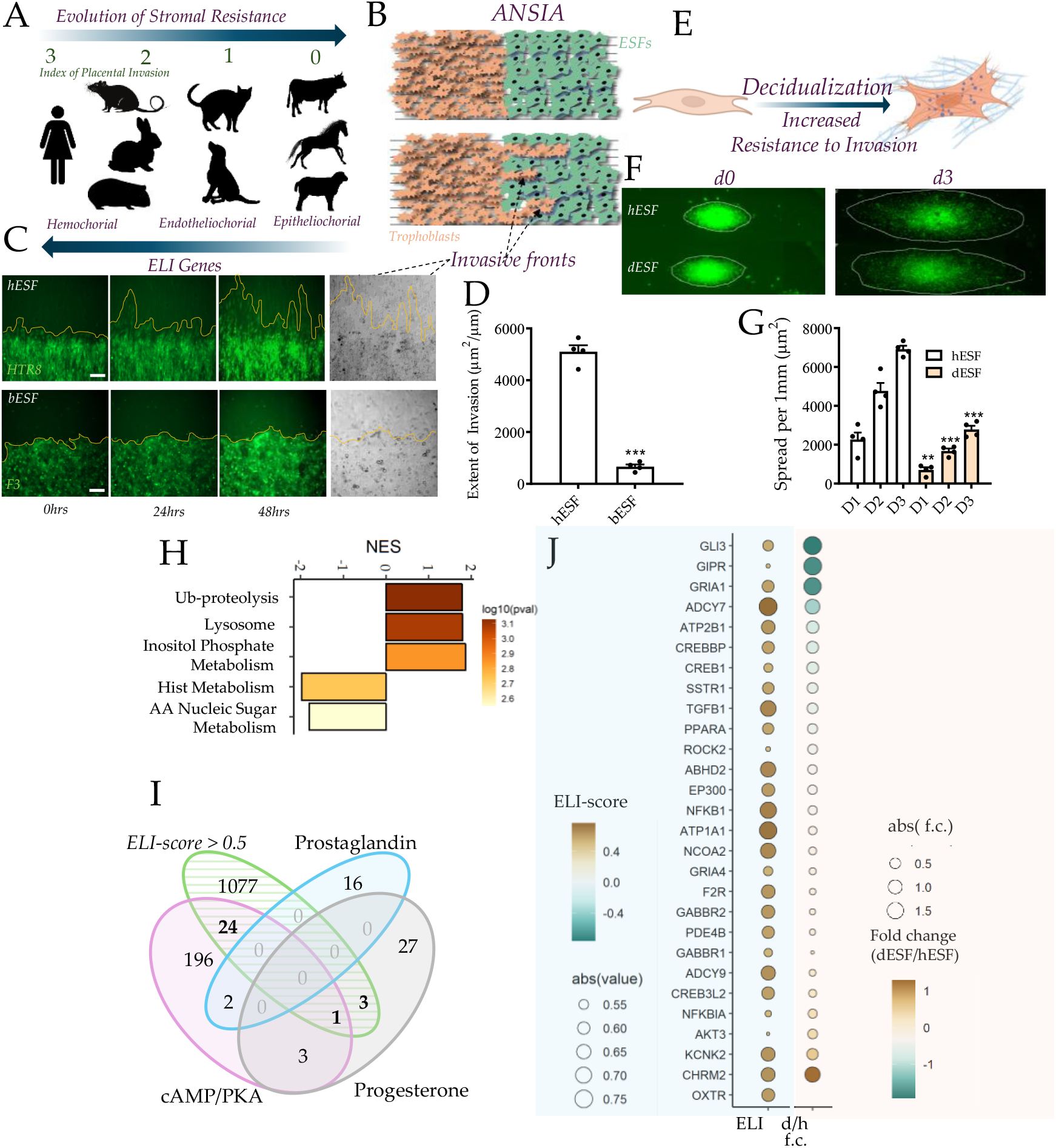
Identification of evolved stromal genes contributing to vulnerability to placental and cancer invasion. **(A-G) Epitheliochorial and decidual ESFs in hemochorial species are characterized by high resistance to trophoblast invasion. (A)** ESFs from species in our analysis to identify ELI genes in stroma correlating with placental invasion; (B) ANSIA quantitatively measures stromal invasion of trophoblasts, with (C) stills from invasion of HTR8 into hESFs, and F3 into bESFs, (D) Quantification. (E) ESF decidualization; (F) HTR8 spheroid invasion on anisotropic hESF, and decidualized hESFs (dESFs) monolayers, quantified in (G**). (H-J) Limited evolved stromal vulnerability gene-set:ELI-D1:** (H) Pathway enrichment in ELI genes; (I) Venn diagram showing overlap of key decidual pathways and gene counts with ELI score> 0.5; (J) ELI-score (left), and fold-change in decidualization for ELI-D1 genes.

We had previously identified ELI genes by differential transcriptomic analysis of ESFs from nine different mammalian species in correlation with a placentation invasion index^3^. Gene-set enrichment revealed pathways related to proteasomal or lysosomal proteolysis, and metabolism, which although interesting, has an unclear connection to stromal resistance (**Fig. 1H**). We hypothesized that parallel accomplishment of similar stromal resistance in (i) evolution of epitheliochorial placentation, and (ii) decidualization of ESFs may converge on similar pathways. We therefore looked for significant ELI genes (ELI score > 0.5) that are part of the pathways critical for decidualization: Prostaglandin biosynthesis, cAMP/Protein Kinase A signaling, and Progesterone signaling (**Fig. 1I**). Among the three decidual gene sets, 28 genes overlap with ELI genes, 24 of them are regulators of PKA signaling. We call this gene-set ELI-D1 (ELI-Decidual Gene Set 1) and found that they also showed reduced expression in hESF post decidualization (**Fig. 1J**).

### Human cancers negatively enrich ELI-D1 genes in stage transition preceding metastasis

We bioinformatically tested the hypothesis that ELI-D1 gene-set is linked to stromal vulnerability to invasion using stage-wise stratified data in The Cancer Genome Atlas (TCGA). GSEA for ELI-D1 genes across stages for many solid cancers showed significant negative enrichment, particularly in the T1 to T2 transition, which characterizes growth of tumor size, frequently preceding onset of dissemination. For most solid TCGA cancers, ELI-D1 was negatively enriched in the T1->T2 transition suggesting that the putative pro-invasable effect of ELI-D1 precedes cancer dissemination. The leading edge genes for various cancers were ABHD2, ADCY7, ADCY9, AKT3, ATP1A1, ATP2B1, CHRM2, CREB1, CREB3L2, CREBBP, EP300, F2R, GABBR1, GIPR, GLI3, GRIA1, NCOA2, NFKB1, PPARA, SSTR1, TGFB1 (**Fig. 2B**). Pancreatic adenocarcinoma (PRAD) uniquely showed an opposite trend, wherein the T1->T2 transition, as defined in TCGA annotation, was positively enriched. To test deeper, we used previously published scRNAseq data of 27 PRAD patients ^7^, and computed the ELI-D1 gene enrichment only within CAFs during stage transitions. We found that ELI-D1 genes were negatively enriched in stage II->IIA, and then positively enriched for IIA->IIB. It is the latter transition, IIA->IIB in PRAD, which is associated with cancer dissemination to proximal lymph nodes, showing a consistent pattern across multiple cancer types for negative ELI-D1 enrichment accompanying the transition to a disseminating stage, which eventually leads to stromal trespass and the onset of metastasis (**Fig 2B-C**).

**Figure 2.**
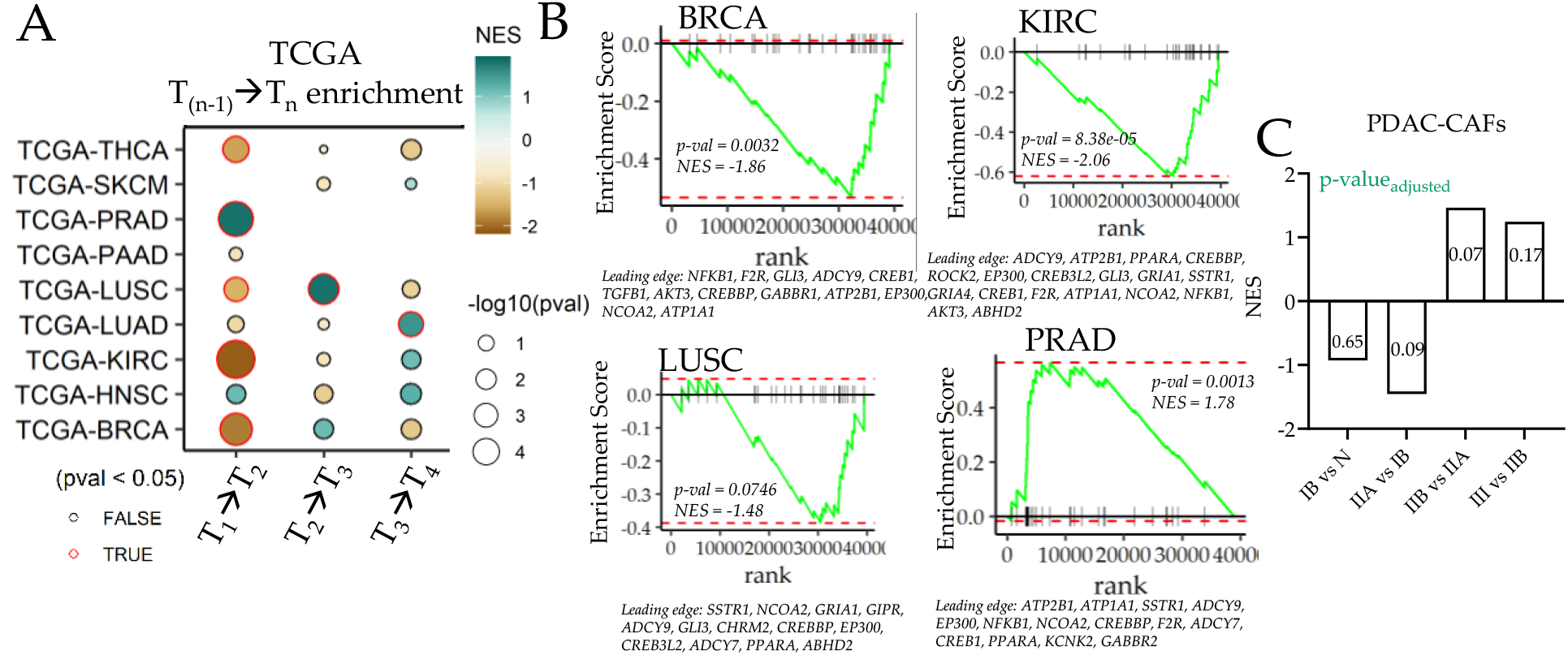
ELI-D1 enrichment in stage transitions in TCGA cancers. **(A)** Enrichment analysis of ELI-D1 in TCGA cancer-types stage transitions. **(B)** GSEA plots showing leading edge ELI-D1 genes in T1->T2 transitions of selected TCGA cancers. **(C)** ELI-D1 enrichment in CAFs from PRAD scRNAseq stage-transitions; p-value in bars.

### Candidate Transcription Factors regulating comparative expression of ELI-D1

Changes in the TF binding sites (TFBS) is a common mechanism of evolution of gene regulation^8^. TFBS changes, both in copy numbers as well as those affecting binding affinity of TFs to the mutated TFBS, can have a dramatic effect in the regulation of downstream gene expression. As a simple linear model, we calculated the regression coefficient β, a measure of the correlation of TFBS copy number in cis-regulatory region of each ELI-D1 gene with its expression ^3^ (**Fig. 3A**).

**Figure 3.**
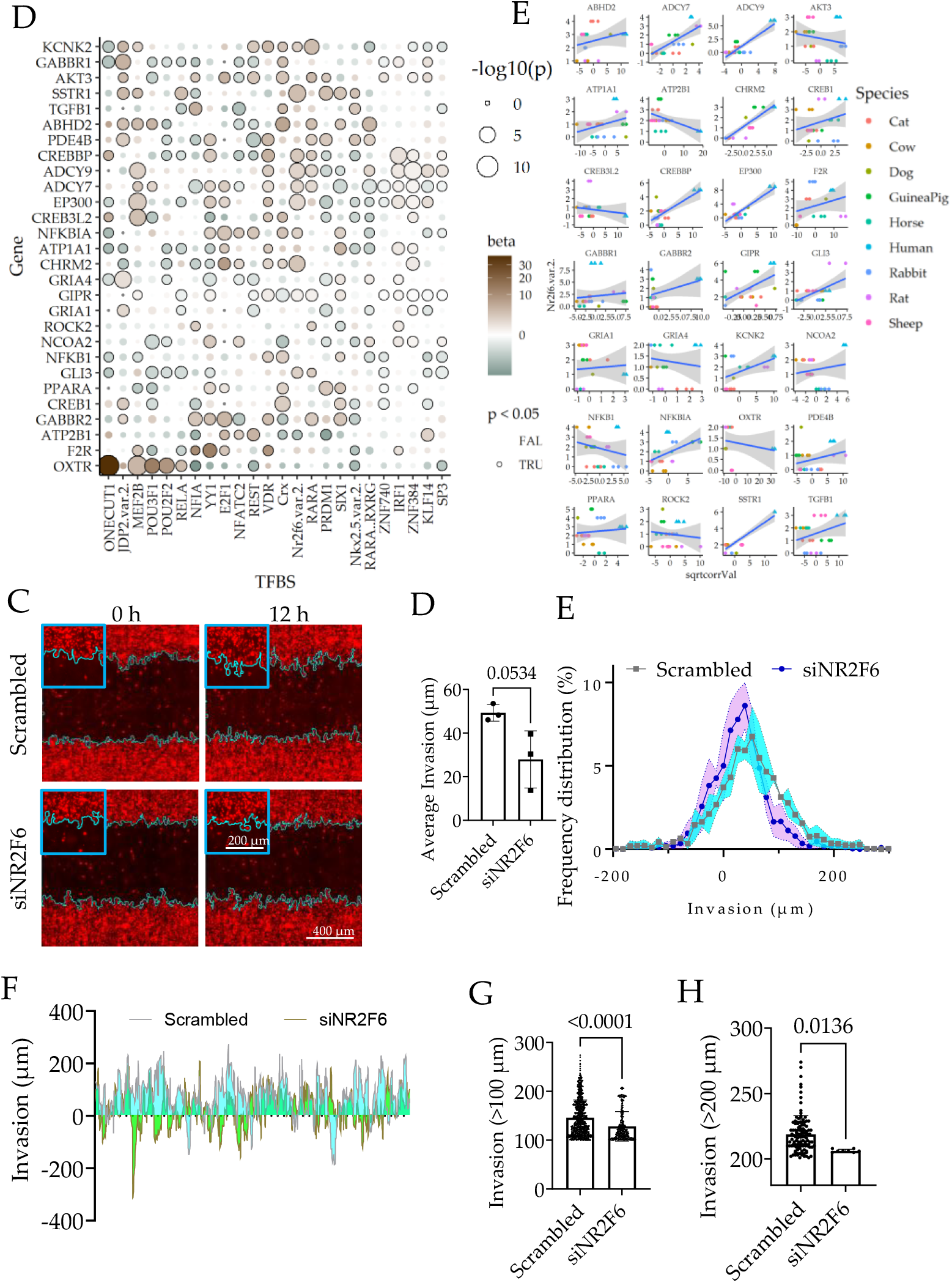
Transcription Factor (TF) candidates regulating ELI-D1 gene expression across mammals. **(A)** TFs β-values explaining ELI-D1 variance across species based on copy number of TF binding sites in cis-regulatory regions of a given gene; **(B)** Example of a selected TF:N2rf26 on variation of gene expression. **(C)** Fluorescent images of A375 (red) invading HDFs (black). Overlayed lines show A375 invading fronts. Inserts showing zoomed in invading fronts. **(D)** Graph showing average A375 invasion depth from three fields of views into scrambled and NR2F6 silenced HDFs. **(E)** Histogram of A375 invasion depth into scrambled and NR2F6 silenced HDFs. **(F)** A375 invasion depth across different horizontal positions in the field of view. **(G-H)** Graph showing invasion depth deeper than 100 μm and 200 μm. For scrambled and siNR2F6, N=1092 and 344 in **(G)**; N= 112 and 8 in **(H)**, respectively.

While there was no overarching single TF regulating comparative ELI-D1 expression, many significant ones informed expression of many leading-edge genes.

Our analysis showed that more than one TF likely played a role in regulating the differential expression of stromal genes across mammals regulating invasion. We chose Nr2f6, which had a broad and significant effect on a large subset of ELI-D1 gene expression across species (**Fig. 3B**), and tested its role in regulating stromal invasion. Towards this, we used ANSIA to measure the effect of NR2F6 gene silencing on human skin fibroblasts resistance to invasion of A375, a cell line derived from malignant melanoma. NR2F6 silencing resulted in significantly reduced aerial invasion (**Fig. 3C-D**). Frequency distribution of invasive forks into hESF monolayer showed a reduction in deep invasive forks (**Fig. 3E-F**). Comparison of deeply invasive forks (>100 um) and (>200 um) showed that NR2F6KD cells were significantly more resistant to deep invasion (**Fig. 3G-H**).

## Discussion

We had previously reported that epitheliochorial placentation is primarily due to the evolution of stromal resistance to invasion, correlating with decreased cancer malignancy^2^. Using ESFs from 9 species, we have also identified stromal genes correlating with their invasibility, termed ELI genes^3,9^. As differential expression is not sufficient to link genes to phenotypes in distant species, we hypothesized that decidualization induced resistance in ESFs may use similar pathways. The identified ELI-D1 gene-set mostly contains inhibitors of cAMP/PKA pathway, not obvious candidates based on gene prioritization. Being limited, they can be easily validated experimentally without the need of a screen. As a means of translational validation, we found that ELI-D1 gene set is negatively enriched for many human cancers preceding onset of their transition to metastasis. CAFs are known to mount a resistive response to growing tumor, which when fails to resist, permits cancer to trespass stroma. ELI-D1 genes are significantly enriched across many cancers in this transition. Finally, we also found several putative candidate transcriptional regulators of evolved stromal vulnerability. We also functionally showed that Nr2f6 may contribute to differential expression of many ELI-D1 genes, and may contribute to increased stromal vulnerability to placental, and cancer malignancy in hemochorial vs epitheliochorial species.

Stromal contribution to placental and cancer invasion is widely appreciated, but search for underlying causal genes is still on, primarily as genotype-phenotype correlation necessitates large data, yet unavailable. Our ELI hypothesis explaining evolution of stromal resistance at MFI, a site of high genetic selectivity and phenotypic variance across mammals has revealed genes that contribute to the inception of cancer dissemination, opening opportunities for anti-metastatic stroma-targeted therapeutics.

## Methods

### Bioinformatics

ELI scores were assigned to genes based on previously described method^3,9^. Pathway enrichment was computed using Gene Set Enrichment Analysis^10^. Cancer patient population gene expression data and staging was downloaded from TCGA^11^, and gene set enrichment analysis performed for each transition stages^10^. To count transcription factor binding sites, the genomic region 5kb upstream to 1 kb downstream of the translation start site for each gene was searched using the FIMO package^12^ with default parameters. RNAseq data for decidualization is taken from Suahil, Y. et al^5^.

### Stromal Invasion Assay

Human dermal fibroblasts (HDFs, PCS-201-012, ATCC) were seeded at 60% confluency. NR2F6 gene was silenced using siRNA (IDT) with RNAiMAX (Thermo Fisher) following the manufacturer’s protocol for two days. A375 cells are labeled with Vybrant DiI (Thermo Fisher) and seeded in 96-well plate with nanopatterns at 40,000 cells/well. A cell free area in each well was created using BioTek AutoScratch (Agilent). HDFs were then detached and seeded into the scratched 96-wells (three wells per condition). Time-lapse images of A375 invading HDFs were taken using Zeiss Oberserver Z1 microscope. A375 invasion profile was analyzed using Fiji/ImageJ with wound healing size tool^13^. 2999 separate widths from each well were analyzed at time 0, and 24h.

## Acknowledgments

We are grateful to our funders: NCI (5U54CA209992; PI: GW, co-I: K), NICHD (K99HD105973; PI: YS), and NICHD (1R01HD112424 ;PI:K), and startup by UConn Health BME.

## Author Contribution

Y.S. conceptualized and performed all bioinformatics analysis, and modeling; W.D. performed functional ANSIA experiments; G.W. provided co-supervision of efforts, funding, and conceptualization of ELI; J.A. contributed to ELI-D1 identification and writing; K. supervised manuscript, funding, conceptualization and wrote the draft.

## Data Availability

All raw data in the figures will be made available on request. Sequencing data are available in NCBI GEO: (GSE197810 ID:200197810), and BioProject (PRJNA564062).

